# Rheology of Growing Axons

**DOI:** 10.1101/2022.04.01.485819

**Authors:** Hadrien Oliveri, Rijk de Rooij, Ellen Kuhl, Alain Goriely

## Abstract

The growth of axons is a key process in neural system development, which relies upon a subtle balance between external mechanical forces and remodeling of cellular constituents. A key problem in the biophysics of axons is therefore to understand the overall response of the axon under stretch, which is often modeled phenomenologically using morphoelastic or viscoelastic models. Here, we develop a microscopic mixture model of growth and remodeling based on protein turnover and damage to obtain the macroscopic rheology of axonal shafts. First, we provide an estimate for the instantaneous elastic response of axons. Second, we demonstrate that under moderate traction velocities, axons behave like a viscoelastic Maxwell material. Third, for larger velocities, we show that failure takes place due to extensive damage.

---

Neurological functions rely on the exchange of electrochemical signals between neurons via slender cellular processes called *axons* [1, 2]. During early neurodevelopment, neuronal cell bodies project axons that extend through the extracellular environment to connect with other target cells [3, 4]. Then, once connected via synapses, axons passively elongate to accommodate the growth of the embedding medium [5, 6]. During this so-called *stretch growth* phase, growth kinematics is fully dictated by the animal’s body expansion. In normal growth conditions, axonal elongation is supported by the addition of cell material, allowing the axon to sustain stretch and maintain structural homeostasis [7–10]. However, upon faster stretch, this mechanism may fail, triggering a cascade of pathophysiological responses that, ultimately, converge to irreversible axonal damage [11–16]. A question is then: How does the axon respond mechanically and structurally to various stretch rates?

Typically, macroscopic – viscoelastic or morphoelastic – models are used to capture the mechanical response of growing axons [7, 12, 17–25]. These simple models can be easily treated mathematically and compared with experiments [26]; however, they are phenomenological and are not explicitly linked to the microstructural changes occurring in the axoplasm during growth. Alternatively, detailed computational models have been proposed to study the role of individual proteins within the cytoplasm [13, 27–39]; typically, a core of parallel *microtubules* cross-linked by microtubule-associated *tau proteins* [Fig. 1(a)]. This approach captures the subtle mechanical interactions between key molecular actors, and the emergent rheology; however, it is relatively complex and does not produce macroscopic models.

**Figure 1.**
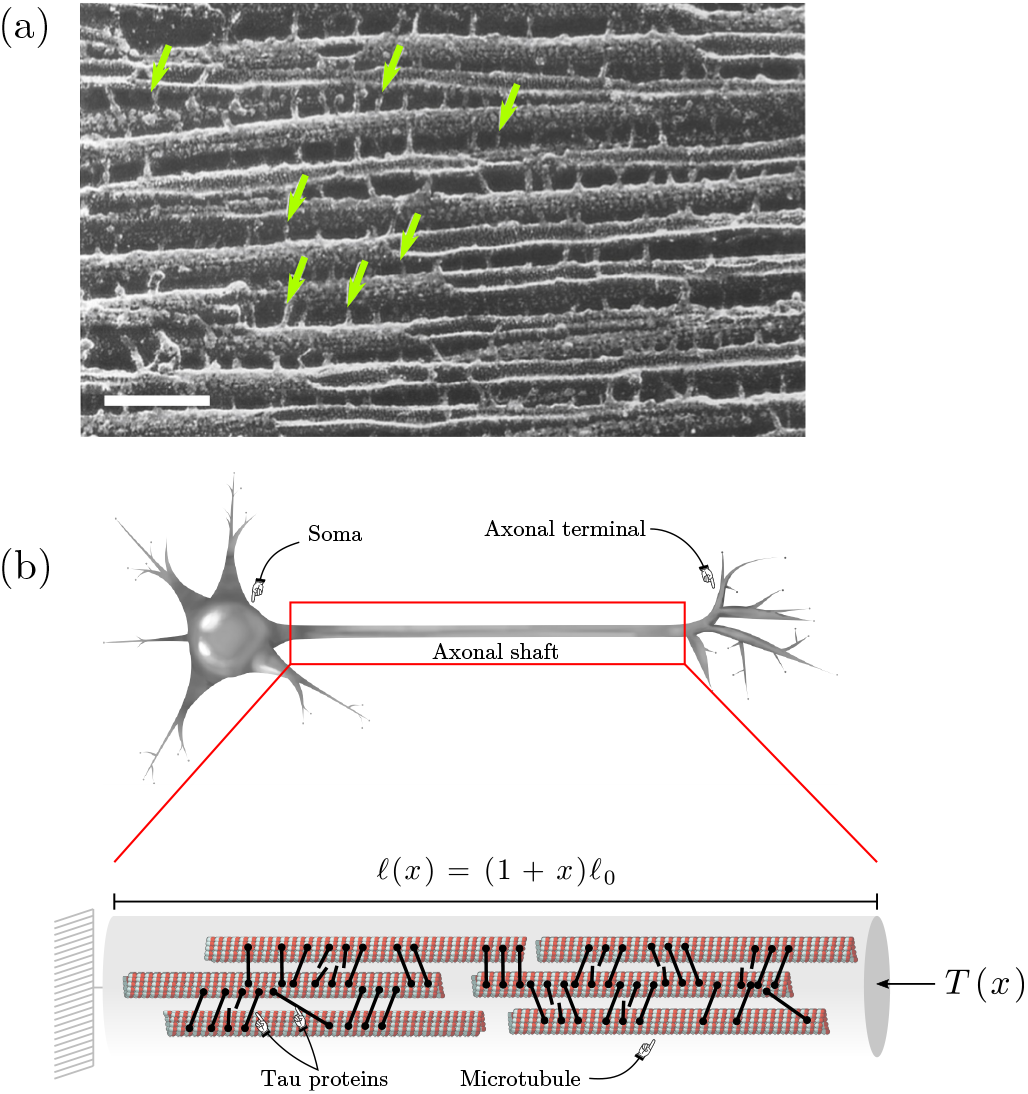
(a) Electron microscopy image of porcine brain microtubules cross-linked with tau proteins shown by arrows. Scale bar = 100 nm. Adapted from [41]. (b) Neuronal axon subject to an imposed strain *x* (*t*). The cytoskeleton is modeled as a well-mixed phase composed of microtubules and tau proteins that may attach, and then detach depending on their individual tension *F*. The individual protein tensions results at global scale in a macroscopic tension *T* (*x*).

Here, we start with the key structural constituents of an axon, microtubules and tau, to derive a macroscopic, homogenized rheological model of axonal stretch growth. We show that, under moderate pulling velocities, the axon behaves like a viscoelastic Maxwell material [24, 40], and the model captures the stress-strain response predicted by previous computational approach [32]. Conversely, for higher stretch rates, remodeling is not fast enough and cross-linking becomes deficient.

## Model

Our model contains two main structural elements: the microtubules and the cross-linking tau proteins. We consider initially a homogeneous cylindrical axon of initial length *ℓ*_0_ with *M*_0_ parallel microtubules of length *a* cross-linked through *N*_0_ tau proteins [Fig. 1(b)]. We study a steady traction scenario in which the axon is towed at a constant stretch rate *ξ*, so that its current length is given by *ℓ* (*t*) = *ℓ*_0_ (1 + *ξt*). Assuming that the cytoplasmic constituents form a well-mixed phase, and neglecting inertial effects, the strain *x* = *ξt* is uniform along the axon [as seen in the kymograph shown in 31]. As the axon elongates, microtubules slide with respect to one another, which stretches the cross-links and promotes their detachment. Following [13, 32, 36, 42], we assume that the tau dissociation kinetics follows a Bell-type model [43] in which a population of *N* (*t*) cross-links subject to a force *F* detaches according to

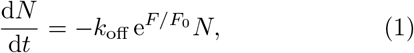

where *k*_off_ is the load-free dissociation rate; and *F*_0_ is a characteristic bond force. We model each protein as a linear spring with constant *κ* and deformation *δ*, which provides the force *F* = *κδ*. Since microtubules are orders of magnitude stiffer than tau proteins [44–47], we further postulate that they remain rigid.

A difficulty is that cross-links attach to different microtubules, that may slide with respect to one another with different velocities [48]. In addition, in a remodeling axon, different cross-links are formed at different times, which requires modeling the axoplasm as a mixture of cross-links with different mechanical states. Assuming a protein is attached at strain *x′*, its deformation at *x* ≥ *x′* is modeled as

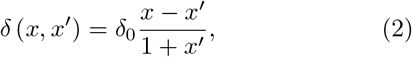

where 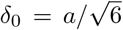 is obtained by a strain-energy based homogenization argument [48]. The total population of cross-links at strain *x* is then given by the mixture [42]

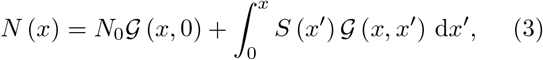

where *S* is the binding rate (per unit strain *x*); and where the kernel,

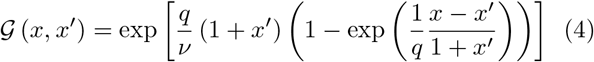

is obtained by solving (1) in terms of *x* for some initial strain *x′*. The dimensionless parameters *q* := *F*_0_*/κδ*_0_ and *ν* := *ξ/k*_off_ characterize the bond dissociation force and the pulling speed, respectively. The first term in (3) represents the decaying population of initial cross-links. The second term accounts for the cross-links formed at all strains 0 ≤ *x′* ≤ *x* during traction, and disconnecting progressively as *x* increases. To simplify notations, we introduce a mixture operator 𝔐 that can be applied to any extensive property 𝒫 (*x, x′*) of the proteins as

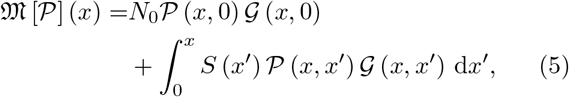

so that, e.g., (3) becomes *N* = 𝔐 [1].

The mechanical response of the axon under applied stretch can be deduced from the individual cross-link strain energies,

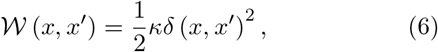

from which we obtain the total energy of the system, *W* = 𝔐 [𝒲]. By the principle of virtual work for an axon under tension *T* (*x*), we have *Tδℓ* = *Tℓ*_0_*δx* = *δW*, where the virtual work *δW* is the sum of all virtual works due to the cross-links:

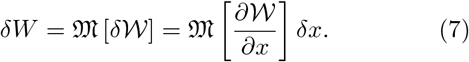

Thus,

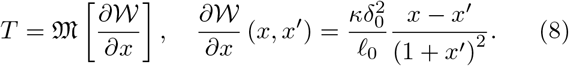

The state of the mixture depends on the attachment of new cross-links with rate *S* (*x*) from either new free tau proteins supplied by the cell, or detached proteins that can form new connections [32, 37]. Assuming a pool of *Ñ* available proteins with uniform concentration *Ñ /ℓ* along the axon (non-limiting transport), a simple model for *S* is *S*d*x* = *k*_on_*Ñ*d*t*, where *k*_on_ is an effective on-rate constant [49]. The number of free proteins then follows

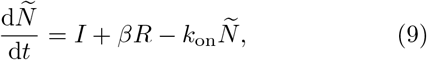

where *I* is a source term; *R* is the number of cross-links disconnected per unit time (1, 2):

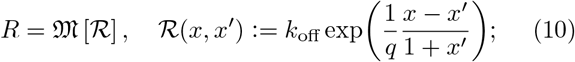

and *β* is the probability of a protein being available for reattachment after disconnection. We hypothesize that the cell aims to maintain a target lineal density *Ñ*_0_*/ℓ*_0_ (i.e. number of cross-links per unit longitudinal length), therefore we posit *I* = *I*_0_(*Ñ*_0_(1 + *x*) − *Ñ*), with *I*_0_ a constant.

The case *β >* 0 is solved numerically [48]. Figure 2(a) shows the effect of reattachment on the primary cross-link population – the population initially present in the axon – in the absence of synthesis (*I*_0_ = 0). As expected, for *β <* 1, the cross-links are eliminated faster than exponentially; larger *β* promotes slower elimination, as cross-links can operate longer. For *β* = 1 however, the cross-links disconnect and reattach endlessly and *N* →*N*_0_*K/* (1 + *K*), with *K* := *k*_on_*/k*_off_ the binding constant.

**Figure 2.**
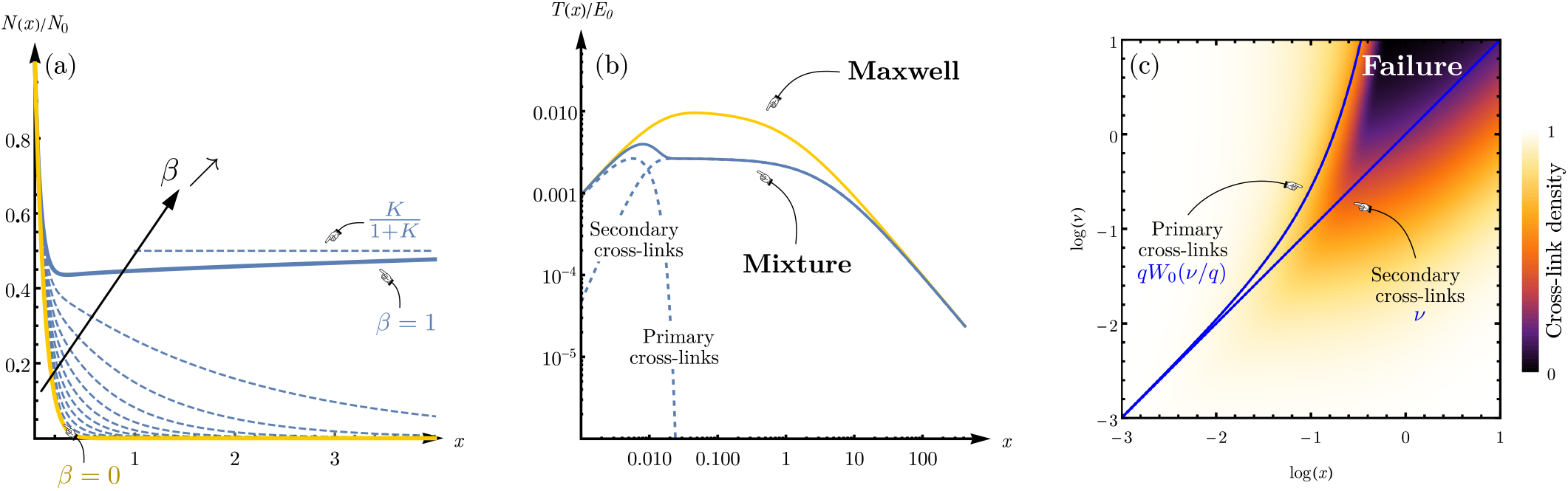
(a) Effect of protein reattachment *β* for a initial population of cross-links (9, 10) simulated numerically for different values of *β*, and with *I*_0_ = 0 (no synthesis), *k*_off_ = 0.1, *K* = 1, *q* = 1 and *ν* = 0.1 [48]. Blue and red lines respectively show the two extreme cases *β* = 1 (full reintroduction) and *β* = 0 (no reintroduction). (b) Log-log plot of the normalized tension *T/E*_0_ vs. strain *x* (*q* = 0.1, *ν* = 0.01). Yellow and blue solid lines respectively show the Maxwell model (12) and the mixture model. Dashed lines show the contributions of the initial and new cross-links. (c) Density *N* (*x*) */ℓ* (*x*) (normalized by *N*_0_*/ℓ*_0_) vs. strain *x* and parameter *ν* (log scale, *q* = 0.1). Solid and dashed lines respectively show the characteristic length-scales of the initial (*x*_I_) and newly-formed (*x*_II_) populations of cross-links.

We henceforth consider the ideal case where detached cross-links do not reattach, *β* ≈ 0. Assuming *I*_0_ ≫ *k*_on_, a solution to (9) is *Ñ* (*x*) ≈ *Ñ*_0_(1 + *x*) (constant lineal density *Ñ*_0_*/ℓ*_0_), giving

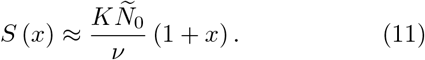

Assuming chemical equilibrium initially, we have *N*_0_ = *KÑ*_0_, and the dynamics is fully governed by *q* and *ν*.

Initially, for small strain *x*, the response under tension due to the primary cross-links is Hookean, namely, *T* (*x*) ≈ *E*_0_*x* with stiffness modulus 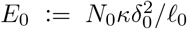 [Fig. 2(b)]. Tension then peaks when *x* = *qW*_0_ (*ν/q*) (where *W*_0_ is the Lambert function). This peak strain provides a typical length-scale *x*_I_ for the primary crosslinks persistence [Fig. 2(c)]. Past the peak, primary cross-links tension quickly vanishes as the cross-links disconnect faster than exponentially (3, 4).

The secondary newly-formed cross-links initially contribute only to ∼ *E*_0_*x*^2^*/*2*ν* to the tension; they do not participate in the linear elastic response, as they are not yet connected and under tension. For large strains however, the total tension is only due to the new cross-links and, as the local strain rate decreases (2), the tension vanishes slowly as *T* (*x*) ∼ *E*_0_*ν/x* [Fig. 2(b)]. Simultaneously, the cross-link density reaches a homeostatic level *N*_0_*I*_0_*/*(*I*_0_ + *k*_on_) ≈ *N*_0_*/ℓ*_0_ with time-scale *x*_II_ = *ν* [Fig. 2(c)].

Depending on whether stretch is applied in a physiological, experimental or traumatic context, the parameter *ν* may be either very small or very large. For rapid stretch, *ν* ≫ 1, we see that *x*_I_ ≪ *x*_II_: The primary crosslinks disconnect before the secondary cross-link density reaches a sufficient level to maintain integrity, and the core ruptures [Fig. 2(a, b)]. Note that, in this regime where density decreases, the well-mixedness assumption fails as random heterogeneities and inertial effects dominate, and the number of loadpaths along the axon also decreases [13, 31]. Conversely, for slow towing, *ν* ≪ 1, we have *x*_I_ ≈ *x*_II_: New cross-links replace the disconnected ones and rescue the axon core. The critical tau deficit, at which the axon is most vulnerable, is given by *d* = max_*x*_ {1 − *N* (*x*) */N*_0_ (1 + *x*)} and, for *ν* small, we have *d* ≈ (1 + *q*^*−*1^)*ν*. Remarkably, in this regime, the model reduces to a viscoelastic Maxwell-like material, with extensional stiffness *E*_0_ and effective viscosity *η*_0_ := *E*_0_*/k*_off_ [48], namely:

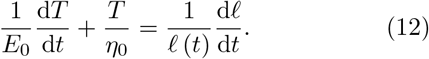

Note that this is not the standard form a Maxwell material which has non zero tension at infinity. Here, our Maxwell-like material has the property that tension goes to zero at infinity as every point grows.

For an axon of diameter ∼0.5 µm [32], with *N*_0_*/ ℓ* _0_ ≈ 100 µm^-1^ [32], *κ* = 0.01–0.1 pN.nm^-1^ [37, 48], and *a* = 10 µm [32], we estimate a Young’s modulus of order ∼10– 100 kPa, which compares with the value reported by [19]. Combining this estimate with measured axon viscosity *η*_0_ = 10^6^–10^7^ Pa.s [21], we estimate *k*_off_ ≈ 10^*−*3^–10^*−*1^ s^-1^, however considering reattachment should in principle yield larger estimates for *k*_off_. The force *F*_0_ can be expressed as *F*_0_ = *k*_B_*T/χ* with *χ* the typical bond separation distance; and *k*_B_*T* ≈ 4 pN.nm. Then, estimating *χ* ≈ 1 nm we obtain *F*_0_ ≈ 1–10 pN [31] and *q* ≈ 10^*−*3^– 10^*−*1^.

## Discrete simulations

Last, we compare our homogenized model against the discrete finite-element model detailed in [32]. We consider a bundle of *M*_0_ ≈ 50 randomly placed parallel microtubules, connected via *N*_0_ ≈ 5000 dynamically breaking cross-links, and we test four different pulling velocities *ν* = 0.01, 0.04, 0.07 and 0.1. For simplicity, we here ignore remodeling (*β* = 0, *S* = 0) [see 32, 48, for details]. We see in Fig. 3 that for moderate velocities, the homogenized model reproduces the stress-strain curves from the discrete simulations faithfully; in particular, note the excellent approximation in the initial Hookean regime (*x* ≪ 1), obtained with no fitting parameter. For higher stretch rates however, as expected, localized heterogeneities dominate the yielding process, which results in our model overestimating the tension.

**Figure 3.**
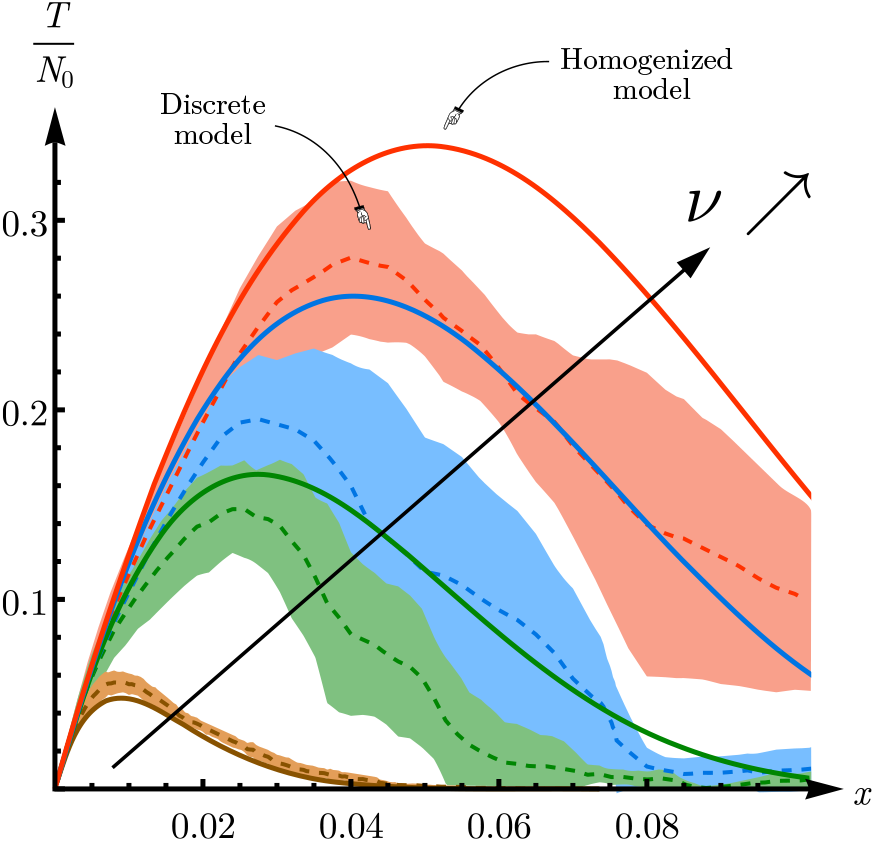
Comparison of our homogenized model with the discrete finite-element model of [32]. Tension-strain curves *T/N*_0_ vs. *x*, for *ν* = 0.01, 0.04, 0.07 and 0.1. Solid lines correspond to the homogenized model. Colored streaks show mean and standard deviation for five different simulations of the discrete model [48].

## Discussion

Healthy growth relies on a subtle balance between mechanics, transport and synthesis of proteins, and cell remodeling [24]. Using mixture theory combined with a Bell-type model for tau disconnection [13, 32, 36, 42, 43], we propose a mechanistic macroscopic model for the mechanical response of stretched axons. In contrast to more realistic, but mathematically intractable, computational models [13, 27–37], this coarsegrained approach establishes a direct mathematical link between cellular parameters and the emergent rheology of axons. First, we derived an expression of the axon’s extensional stiffness: *E*_0_ ≈ *N*_0_*κa*^2^*/*6*ℓ*_0_. Denoting *ρ* and 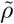 the lineal densities of cross-linking and microtubules, respectively, we obtain the scaling law 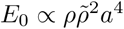, that explicitly relates the microstructural geometry to the overall elastic response [39]. Second, we showed that rate-dependent effects emerge from the energy-dissipating disconnection of linearly elastic cross-links embedded in a dynamically-evolving mixture, as shown also in [31]. For small strain rates, we proved that the system behaves like a Maxwell viscoelastic material, with extensional stiffness *E*_0_ and viscosity *η*_0_ = *E*_0_*/k*_off_. This prediction recovers the observed fluid-like behavior of axons [12, 18, 19, 21, 22] and corroborates previous computational studies [31]. For higher strain rates however, a critical regime appears where the axon fails to maintain a sufficient level of cross-linking due to insufficient material synthesis, with a critical tau deficit attained around the characteristic strain *x*_I_ = *qW*_0_ (*ν/q*) ≈ *ν*. Note, however, that subtler rate-dependent effects could potentially emerge from more sophisticated viscoelastic models of individual tau and microtubules [36, 37], or by taking into account the actomyosin sheath that generate active tension [22, 33, 50]. Neuronal injury involves many other mechanisms such as microtubule breakage and collapse [15, 16, 36, 37, 51]. Here, we have ignored these effects of extreme axonal mechanics to focus our attention on the evolution of microtubule cross-linking during slow growth. By definition, growth is limited by mass uptake [24], and, in singularly large and fast growing cells like neurons, an important question is what mechanisms regulate material availability [5, 9]. Here, we found that a linear active coupling between axonal length *R* and protein synthesis rate *I* was adequate to maintain sufficient material supply. Biologically, this modeling assumption implies that the cell is able to sense its current length to regulate the production of new proteins, through a mechanism that is not yet understood [10, 52].

Axonal growth is the central components of neurodevelopment. It obeys intricate rules with multiple interplays between mechanics, kinematics, biological feedback, remodeling and protein supply. The detailed response of axons to stimuli and loads remains elusive, yet, based on universal microscopic principles related to attachment and detachment of cross-links, their macroscopic response can be obtained and systematically compared with experiments by targeting specific microscopic properties. This type of approaches opens the door to more refined multiscale theories of axonal growth.

A. G. acknowledges support from the Engineering and Physical Sciences Research Council of Great Britain under research Grant No. EP/R020205/1. E. K. acknowledges support from the National Science Foundation under research Grant CMMI 1727268. The authors thank Kristian Franze and Kyle Miller for insightful discussions.

## Notes

### Competing Interest Statement

The authors have declared no competing interest.

