## Supplementary Material for "Rheology of Growing Axons"

### Supplemental Material for *Rheology of Growing Axons*

Hadrien Oliveri[1], Rijk de Rooij[2], Ellen Kuhl[2] and Alain Goriely[1*]
[1]*Mathematical Institute, University of Oxford, Oxford, OX2 6GG, UK and*
[2]*Department of Mechanical Engineering, Stanford University, Stanford, CA, 94305, USA*

(Dated: April 1, 2022)

#### I. DETAILS OF THE MODEL

Here, we provide details on the assumptions underlying the mixture model developed in *Main text*. We consider the two main structural elements of the axon, i.e. rigid microtubules cross-linked by stretchable tau proteins. Microtubules are long filament-like polymers that, in the axon, are mostly aligned longitudinally and parallel to one another. We assume that they are fully aligned to the longitudinal axis of the axon, and that they can only move by translation along that axis of the axon. Hence, our model is one-dimensional and we are interested in longitudinal mechanical response of the axon as a result of the interaction between microtubules and tau proteins. [1–9].

##### A. Kinematics of tau protein deformation

Here, we study how a given cross-link deforms when the whole axons undergoes a strain $x$. In a deformation of the axon, microtubules slide with respect to one another, which in turn stretches the connecting cross-links. For simplicity, we neglect the growth and shrinkage of microtubules that might result in net migration of the microtubules along the axon. Thus, we assume that all microtubules have identical and constant length $a$. Considering that microtubule cross-links are homogeneously distributed along the axon, we also assume that each microtubule's center of mass (midpoint) moves according to an affine transformation, i.e. its center of mass is passively advected with the embedding medium, so that the longitudinal motions of microtubules are imposed geometrically. Namely, if $p$ is the longitudinal position of a microtubule's midpoint, then for any strains $x$ and $x'$ of the axon, we have:

$$\frac{p(x)}{1+x} = \frac{p(x')}{1+x'}. \tag{S1}$$

Now, consider a cross-link created at strain $x'$, and connected to two different microtubules $i$ and $j$ with respective positions $p_i$ and $p_j$. Neglecting the nonlinear effects due to the small lateral distance between the two microtubules, the gradient of tau displacement with respect to the global strain $x$ is given by

$$\frac{\partial \delta_{ij}}{\partial x} = \left| \frac{\partial p_j}{\partial x} - \frac{\partial p_i}{\partial x} \right| = \frac{|p_j(x') - p_i(x')|}{1+x'}, \tag{S2}$$

which can be integrated to obtain

$$\delta_{ij}(x, x') = |p_j(x') - p_i(x')| \times \frac{x - x'}{1+x'}. \tag{S3}$$

Hence, different cross-links attached at the same strain $x'$ will undergo different deformations, depending on the relative positions of their microtubules (Fig. S1).

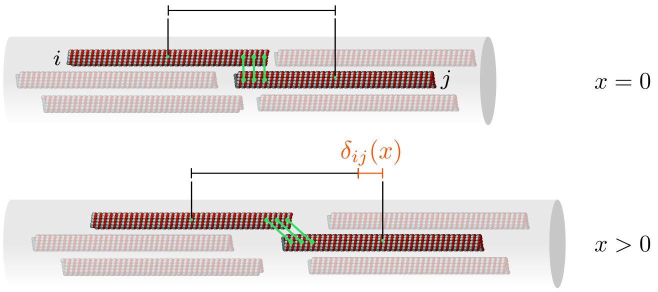


FIG. S1. Deformation of cross-links connecting two microtubules $i$ and $j$. The rate of deformation depends on the distance between the two microtubules, whose center of mass undergoes affine displacement.

##### B. Elastic response

Here we focus on the initial Hookean elastic response, with $x' = 0$ and $x \ll 1$, and we assume that the cross-links do not attach or dissociate. The total strain energy of the system can be obtained from (S3) by integrating the individual protein energies with spring constant $\kappa$,

$$\mathcal{W}_{ij}(x) = \frac{1}{2}\kappa \delta_{ij}(x,0)^2 = \frac{1}{2}\kappa (X_j - X_i)^2 x^2, \tag{S4}$$

over all possible initial longitudinal positions $X_i = p_i(0) \in [a/2, \ell_0 - a/2]$ of the $M_0$ microtubules, taken to be uniformly distributed along the axon:

$$W(x)=\left(\frac{M_0}{\ell_0-a}\right)^2\int_{a/2}^{\ell_0-a/2}\int_{a/2}^{\ell_0-a/2}n(X_i,X_j)\,\mathcal{W}_{ij}(x)\,\mathrm{d}X_i\,\mathrm{d}X_j. \tag{S5}$$

Here, $n(X_i,X_j)$ is the number of cross-links connecting two overlapping microtubules located at positions $X_i$ and $X_j$. We assume that $n(X_i,X_j)$ is proportional to the overlapping distance, i.e. $n(X_i,X_j)=\rho(a-|X_i-X_j|)_+$, where $(\cdot)_+$ is the ramp function and $\rho$ is a lineal density of cross-linking linked to $N_0$, the number of cross-links, via

$$N_0=\left(\frac{M_0}{\ell_0-a}\right)^2\int_{a/2}^{\ell_0-a/2}\int_{a/2}^{\ell_0-a/2}n(X_i,X_j)\,\mathrm{d}X_i\,\mathrm{d}X_j. \tag{S6}$$

From (S5), we also obtain the extensional stiffness

$$E_0=\frac{\kappa}{\ell_0}\left(\frac{M_0}{\ell_0-a}\right)^2\int_{a/2}^{\ell_0-a/2}\int_{a/2}^{\ell_0-a/2}n(X_i,X_j)(X_j-X_i)^2\,\mathrm{d}X_i\,\mathrm{d}X_j. \tag{S7}$$

Assuming $a\ll\ell_0$ (long initial axon), (S6) simplifies to

$$N_0=\frac{\rho M_0^2\left(3a^2\ell_0-4a^3\right)}{3(\ell_0-a)^2}\approx\frac{\rho M_0^2a^2}{\ell_0}. \tag{S8}$$

Then we integrate (S5, S7) using (S8), to obtain

$$W(x)\approx\frac{1}{12}N_0\kappa a^2x^2, \tag{S9}$$

and

$$E_0\approx\frac{N_0\kappa a^2}{6\ell_0}. \tag{S10}$$

##### C. Protein dissociation and model simplification

To model the dissociation of proteins we need to track all pairs of microtubules as shown above. Using the kernel provided in *Main text*, (4), (S5, S6) become

$$N(x)=\left(\frac{M_0}{\ell_0-a}\right)^2\int_{a/2}^{\ell_0-a/2}\int_{a/2}^{\ell_0-a/2}n(X_i,X_j)\exp\left[\frac{F_0}{\kappa\nu|X_i-X_j|}\left(1-\exp\left(\frac{\kappa x|X_i-X_j|}{F_0}\right)\right)\right]\mathrm{d}X_i\,\mathrm{d}X_j, \tag{S11}$$

and

$$W(x)=\left(\frac{M_0}{\ell_0-a}\right)^2\int_{a/2}^{\ell_0-a/2}\int_{a/2}^{\ell_0-a/2}n(X_i,X_j)\,\mathcal{W}_{ij}(x)\exp\left[\frac{F_0}{\kappa\nu|X_i-X_j|}\left(1-\exp\left(\frac{\kappa x|X_i-X_j|}{F_0}\right)\right)\right]\mathrm{d}X_i\,\mathrm{d}X_j. \tag{S12}$$

Next, to take into account added proteins, we track all tau associations and microtubule additions occurring at strain $x'\le x$ [*Main text*, (3)], which, ultimately, will result in a triple integration (over $x$, $X_j$ and $X_i$).

To make progress, we instead next posit a characteristic length scale $\delta_0$ and substitute all protein deformations (S3) with a unique deformation

$$\delta(x)\approx\delta_0x, \tag{S13}$$

where $\delta_0$ is chosen so as to obtain the same strain energy given by (S9). The new energy can be easily computed [*Main text*, (6)] and we see that

$$\delta_0=\frac{a}{\sqrt{6}}. \tag{S14}$$

Note that $\delta_0$ is much larger than the typical size of the individual tau proteins [i.e. ~50 nm, while $a\approx10$ μm, see 6], which implies that tau proteins stretch individually much faster than the entire axon does.

We extend this approach to growing axons by assuming that the density of microtubules does not vary much during the growth process, i.e. $M(x)\approx M_0(1+x)$, so that we can use (S13, S14) to represent every internal state of the mixture during the elongation process:

$$\delta\left(x, x'\right) = \frac{a}{\sqrt{6}} \frac{x - x'}{1 + x'}. \tag{S15}$$

Note that this approach is justified by the fact that $\delta_0$ is independent of $\ell_0$, thus (S14) applies in principle to axons of arbitrary size $\ell$.

Fig. S2 shows a comparison of the tension $T\left(x\right)$ and number of cross-links $N\left(x\right)$ obtained from (S11, S12), and from the simplification (S13, S14), respectively. We see that for relatively slow pulling speeds, the approximation is in good agreement with the more detailed model.

#### II. ASYMPTOTICS

We provide details on the asymptotic results given in *Main text*. We first study the case of small and large strains $x \ll 1$ and $x \gg 1$ (Section II A), then we show that for slow stretch $\nu \ll 1$, the tension obtained with our mixture model is given asymptotically by the tension of a Maxwell material (Section II B).

##### A. Cases $x \to 0$ and $x \to \infty$

The case $x \to 0$ is straightforward and can be obtained via a Taylor expansion of the various quantities of interest around $x = 0$. The case $x \to \infty$ can be studied using integration by part. By way of illustration, we derive the asymptotic behavior of the population $N\left(x\right)$ of attached cross-links as $x \to \infty$. For clarity of notations, we can set $N_0 = 1$ and $E_0 = 1$, without loss of generality.

We first perform a change of variable $u = x'/x$ to fix the integration domain:

$$\begin{aligned} N\left(x\right) &\sim \int_0^x S\left(x'\right) \mathcal{G}\left(x, x'\right) \mathrm{d}x' \\ &= x \int_0^1 S\left(ux\right) \mathcal{G}\left(x, ux\right) \mathrm{d}u \\ &= x \int_0^1 S\left(ux\right) \mathrm{e}^{g(u)} \mathrm{d}u, \end{aligned} \tag{S16}$$

with

$$g\left(u\right) = \frac{q}{\nu}\left(1 + ux\right)\left(1 - \exp\left(\frac{x}{q}\frac{1-u}{1+xu}\right)\right). \tag{S17}$$

We then extract the leading order through integration by part:

$$\begin{aligned} N\left(x\right) &\sim x \int_0^1 S\left(ux\right) \mathrm{e}^{g(u)} \mathrm{d}u \\ &= x \int_0^1 \frac{S\left(ux\right)}{g'\left(u\right)} \mathrm{d}\left(\mathrm{e}^{g(u)}\right) \\ &= \nu S\left(x\right) - \underbrace{x \int_0^1 \mathrm{e}^{g(u)} \mathrm{d}\left(\frac{S\left(ux\right)}{g'\left(u\right)}\right)}_{\text{h.o.t.}} \\ &\sim x. \end{aligned} \tag{S18}$$

Similarly, for the tension $T$, we integrate by parts twice to obtain

$$T\left(x\right) \sim \frac{\nu}{x}. \tag{S19}$$

##### B. Case $\nu \ll 1$: Maxwell model

We show that for slow growth $\nu \ll 1$ and fixed $x$, the model reduces asymptotically to the Maxwell model. For clarity of notation, we here note $\varepsilon = \nu$, the small parameter for the asymptotic analysis. In the case of a constant-speed traction considered, the Maxwell model, with extensional stiffness $E_0$ and viscosity $E_0/k_{\text{off}}$, is defined via the tension $\widetilde{T}$ that obeys

$$\widetilde{T}'\left(x\right) + \frac{1}{\varepsilon}\widetilde{T}\left(x\right) = \frac{1}{1+x}, \quad \widetilde{T}(0) = 0, \tag{S20}$$

[*Main text*, (12)], where $(\,\cdot\,)'$ denotes derivative with respect to $x$. The solution to (S20) is

$$\widetilde{T}\left(x\right) = \exp\left(-\frac{1+x}{\varepsilon}\right)\left[\mathrm{Ei}\left(\frac{1+x}{\varepsilon}\right) - \mathrm{Ei}\left(\frac{1}{\varepsilon}\right)\right], \tag{S21}$$

where Ei is the exponential integral function. For small $\varepsilon$, the solution can be expanded to obtain

$$\widetilde{T}\left(x\right) = \frac{\varepsilon}{1+x} - \varepsilon \mathrm{e}^{-x/\varepsilon} + \mathcal{O}\left(\varepsilon^2\right). \tag{S22}$$

Next, we address the asymptotic behavior of the mixture model for $\varepsilon \ll 1$. In this case, the tension is given in terms of the integral

$$\mathcal{I}\left(x\right) = \int_0^x \frac{x - x'}{1 + x'} \mathcal{G}\left(x, x'\right) \mathrm{d}x', \tag{S23}$$

such that

$$T\left(x\right) = x\mathcal{G}\left(x, 0\right) + \frac{1}{\varepsilon}\mathcal{I}\left(x\right). \tag{S24}$$

This integral is of the form

$$\mathcal{I}\left(x\right) = \int_0^x f\left(x'\right) \mathrm{e}^{\lambda g(x')} \mathrm{d}x' \tag{S25}$$

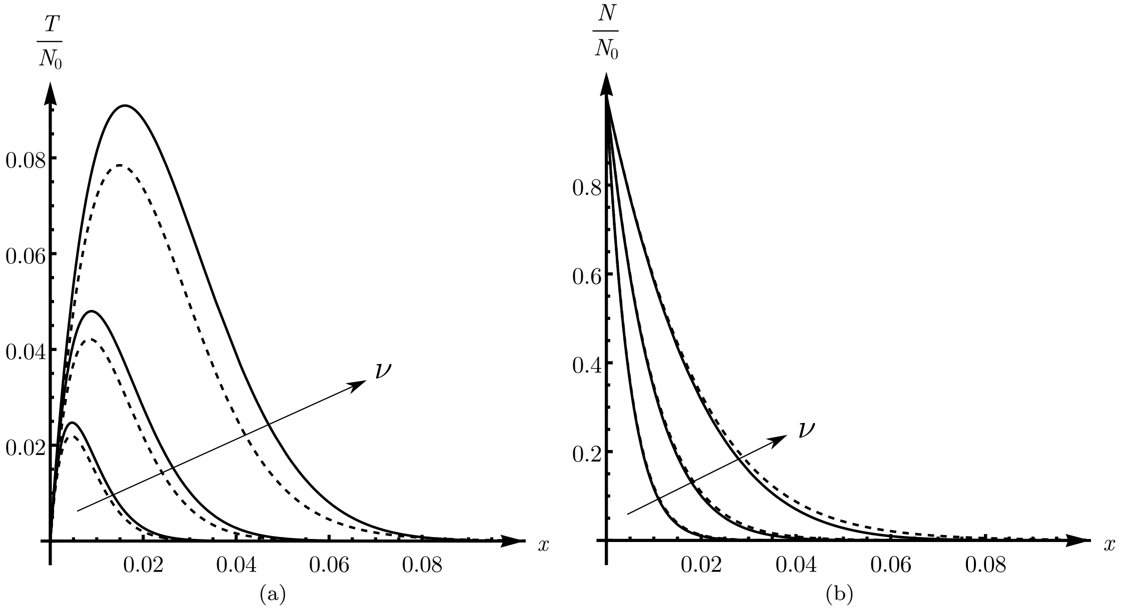


FIG. S2. Axonal tension $T(x)/N_0$ (a) and number of cross-links $N(x)/N_0$ (b) obtained by the integral model (S11, S12) (dashed line), and by the single-lengthscale model (S13, S14) (solid line). Parameters given in Section IV. We use $\nu = 0.005$, 0.01 and 0.02.

with $\lambda = \varepsilon^{-1} \gg 1$; and with $f(x') = (x - x')/(1 + x')$ and $g(x') = q(1 + x')\left(1 - \mathrm{e}^{f(x')/q}\right)$. The argument in the exponential is maximal for $x' = x$, thus, we adapt Laplace's method of integration by expanding $g(x')$ and $f(x')$ to first order around $x' = x$, noting that $f(x) = g(x) = 0$:

$$\begin{aligned}\mathcal{I}(x) &\approx -f'(x)\int_0^x (x - x')\exp\left(-\lambda g'(x)(x - x')\right)\mathrm{d}x' \\ &= \frac{1}{1+x}\int_0^x (x - x')\,\mathrm{e}^{-\lambda(x-x')}\,\mathrm{d}x' \\ &= \frac{1 - (1 + \lambda x)\,\mathrm{e}^{-\lambda x}}{\lambda^2(1+x)} \\ &= \frac{\varepsilon^2 - \varepsilon(\varepsilon + x)\,\mathrm{e}^{-x/\varepsilon}}{1+x}.\end{aligned} \tag{S26}$$

From (S24, S26) we obtain

$$\begin{aligned}T(x) = &\frac{\varepsilon}{1+x} - \frac{x+\varepsilon}{1+x}\mathrm{e}^{-x/\varepsilon} \\ &+ x\exp\left[\frac{q}{\varepsilon}\left(1 - \mathrm{e}^{x/q}\right)\right] + \mathcal{O}\left(\varepsilon^2\right),\end{aligned} \tag{S27}$$

which can be put under the form $T(x) = \widetilde{T}(x)\left(1 + \delta(x) + \mathcal{O}(\varepsilon)\right)$, with

$$\delta(x) = -\frac{x\left((x+1)\exp\left(\frac{q+x}{\varepsilon} - \frac{q}{\varepsilon}\mathrm{e}^{x/q}\right) + \varepsilon - 1\right)}{\varepsilon\left(\mathrm{e}^{x/\varepsilon} - x - 1\right)}. \tag{S28}$$

Clearly, $\delta(x)$ is biggest for $x = \mathcal{O}(\varepsilon)$, therefore, using $X := x/\varepsilon = \mathcal{O}(1)$, we have

$$\delta(X) \approx -\frac{\varepsilon X\left(X^2/2q - X - 1\right)}{\mathrm{e}^X - 1} = \mathcal{O}(\varepsilon), \tag{S29}$$

namely, $T(x) = \widetilde{T}(x)\left(1 + \mathcal{O}(\varepsilon)\right)$, which proves that, to first order, the tension is given by the Maxwell model, see Fig. S3(a). We can also verify numerically that

$$r(x) = \frac{1}{\varepsilon}\left(1 - \frac{T(x)}{\widetilde{T}(x)}\right), \tag{S30}$$

is indeed $\mathcal{O}(1)$, see Fig. S3(b).

#### III. REINTRODUCTION OF CROSS-LINKS

Next, we develop a numerical method to solve the integro-differential problem given by *Main text*, (9). From $S\mathrm{d}x = k_{\mathrm{on}}N\mathrm{d}t$ and $x = \xi t$, we obtain $N = S\xi/k_{\mathrm{on}} = S\nu/K$. Thus the governing equation for $\widetilde{N}$ can be recast so that it only involves $S$, as

$$\frac{\mathrm{d}S}{\mathrm{d}x} + \frac{S}{\nu}\left(K + \frac{I_0}{k_{\mathrm{off}}}\right) = \frac{I_0 N_0}{\nu^2 k_{\mathrm{off}}}(1+x) + \frac{\beta K}{\nu^2}P(x) \tag{S31}$$

with

$$\begin{aligned}P(x) = &\,\mathrm{e}^{x/q}\mathcal{G}(x,0) \\ &+ \int_0^x S(x')\exp\left(\frac{1}{q}\frac{x-x'}{1+x'}\right)\mathcal{G}(x,x')\,\mathrm{d}x'.\end{aligned} \tag{S32}$$

In the case where no new proteins are introduced ($I_0 = 0$), by virtue of the balance of mass, $S$ is bounded and converges to $S_\infty \geq 0$ with

$$S_\infty = \frac{\beta}{\nu}\int_0^x S(x')\exp\left(\frac{1}{q}\frac{x-x'}{1+x'}\right)\mathcal{G}(x,x')\,\mathrm{d}x'. \tag{S33}$$

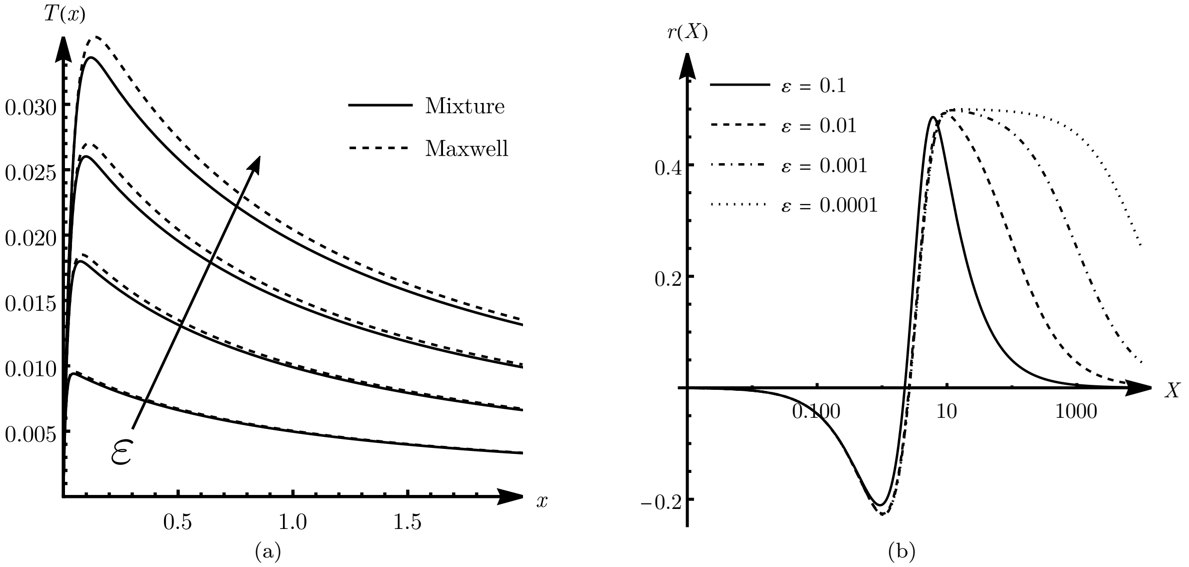


FIG. S3. (a) Tension-strain curve for the Maxwell model (dashed line) and the mixture models (solid line), for $\varepsilon = 0.01$, 0.02, 0.03, 0.04 and $q = 2$. (b) Log plot of the remainder $r(X)$, defined in (S30), vs. $X = \varepsilon x$ for various values of $\varepsilon$ and $q = 2$. We verify that $r(X) = \mathcal{O}(1)$ as $\varepsilon \to 0$.

Using integration by part as in Section II A, we have

$$\int_0^x S(x') \exp\left(\frac{1}{q}\frac{x - x'}{1 + x'}\right) \mathcal{G}(x, x')\, \mathrm{d}x' \sim \nu S_\infty, \quad \text{(S34)}$$

for $x \to \infty$. Thus, unsurprisingly, $S_\infty = 0$ if $\beta < 1$, as the probability of a protein being recycled $n$ times is $\beta^n \to 0$. If $\beta = 1$, proteins are endlessly disconnected and reintroduced, and the number of cross-links stabilizes at $N_\infty = S_\infty \nu = N_0 K/(K+1)$.

The general time-dependent problem (S31) is solved numerically by discretizing the domain as a sequence $x_i = (i-1)h$, $\forall i \in [1, n]$, with $h$ a constant step size; and $n$ the desired number of evaluation points. We discretize (S31) using the backward Euler scheme:

$$\frac{S_i - S_{i-1}}{h} + \alpha_1 S_i = \alpha_2 (1 + (i-1)h) + \alpha_3 P_i, \quad \text{(S35)}$$

with $S_i$ and $P_i$ the estimated values of $S(x_i)$ and $P(x_i)$ respectively; and where

$$\alpha_1 = \frac{1}{\nu}\left(K + \frac{I_0}{k_{\mathrm{off}}}\right), \quad \alpha_2 = \frac{I_0 N_0}{\nu^2 k_{\mathrm{off}}}, \quad \alpha_3 = \frac{K\beta}{\nu^2}, \quad \text{(S36)}$$

are dimensionless parameters. The integral part $P_i$ is computed using the trapezoidal method with step size $h$:

$$\begin{aligned} P_i &= C_{i,1} + \frac{h}{2}\sum_{j=1}^{i-1}\left(C_{i,j+1}S_{j+1} + C_{i,j}S_j\right) \\ &= C_{i,1} + \frac{h}{2}\left(C_{i,1}S_1 + C_{i,i}S_i\right) + h\sum_{j=2}^{i-1} C_{i,j}S_j, \quad \text{(S37)} \end{aligned}$$

where

$$C_{i,j} = \mathcal{G}(x_i, x_j) \exp\left(\frac{1}{q}\frac{x_i - x_j}{1 + x_j}\right), \quad \text{(S38)}$$

are pre-computed coefficients defined for $0 \le j \le i \le n$.

Finally, combining (S35, S37), we obtain the iterative integration algorithm

$$S_i = \mathbb{A}_i + \sum_{k=1}^{i-1} \mathbb{B}_{i,k} S_k, \quad \text{(S39)}$$

where, for all $i \in [1, n]$,

$$\mathbb{A}_i = \frac{h\alpha_3 C_{i,1} + h\alpha_2 (1 + (i-1)h)}{1 + h\alpha_1 - h^2\alpha_3 C_{i,i}/2}, \quad \text{(S40)}$$

and, for all $k \in [1, i-1]$,

$$\mathbb{B}_{i,k} = \begin{cases} \dfrac{h^2\alpha_3 C_{i,1}/2}{1 + h\alpha_1 - h^2\alpha_3 C_{i,i}/2} & \text{if } k = 1, \\ \dfrac{1 + h^2\alpha_3 C_{i,i-1}}{1 + h\alpha_1 - h^2\alpha_3 C_{i,i}/2} & \text{if } k = i-1, \\ \dfrac{h^2\alpha_3 C_{i,k}}{1 + h\alpha_1 - h^2\alpha_3 C_{i,i}/2} & \text{otherwise.} \end{cases} \quad \text{(S41)}$$

Figure S4 shows example computed profiles for $S(x)$, for various values of $\beta \in [0, 1]$ and in the case where no new proteins are introduced ($I_0 = 0$).

#### IV. COMPARISON WITH DISCRETE MODEL

In this section, we adapt our parameters to the particular setting developed in [6]. In this work, the cross-

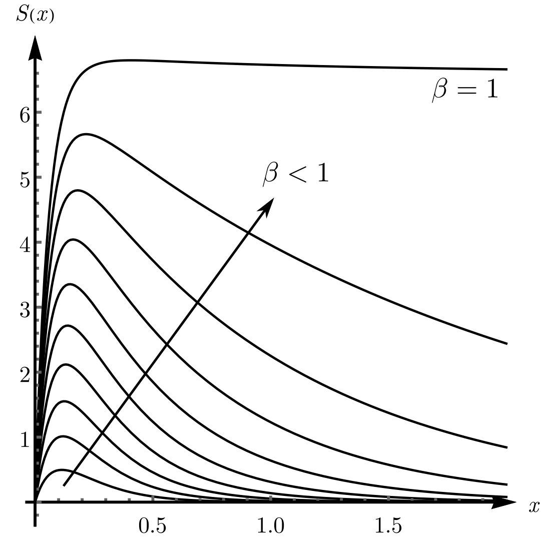


FIG. S4. Reintroduction of disconnected cross-links. Binding rate $S(x)$ vs $x$ for various values of $\beta$. Here we consider the case where no new proteins are synthesized ($I_0 = 0$).

links are modeled as Hookean rods with Young's modulus 10 MPa and cross-sectional area 1 $nm^2$. The cross-link lengths can vary depending on the relative positions of the two anchoring points in the simulation; here, we select the average value 150 nm as reference cross-link length. We deduce the spring constant 0.03 $pN.nm^{-1}$. Note, however, that in the computational model, cross-links are attached with an average angle $\beta \approx 45°$ with respect to the microtubule axis. Thus the equivalent longitudinal spring constant is actually $\kappa = \kappa^* \cos^2 \beta = \kappa^*/2$. All other relevant parameters are identical to those used in [6].

*
